# Predicting Aquatic Development and Mortality Rates of *Aedes Aegypti*

**DOI:** 10.1101/407502

**Authors:** Josef Zapletal, Himanshu Gupta, Madhav Erraguntla, Zach N. Adelman, Kevin M. Myles, Mark A. Lawley

## Abstract

Mosquito-borne pathogens continue to be a significant burden within human populations, with *Aedes aegypti* continuing to spread dengue, chikungunya, and Zika virus throughout the world. Using data from a previously conducted study, a linear regression model was constructed to predict the aquatic development rates based on the average temperature, temperature fluctuation range, and larval density. Additional experiments were conducted with different parameters of average temperature and larval density to validate the model. Using a paired t-test, the model predictions were compared to experimental data and showed that the prediction models were not significantly different for average pupation rate, adult emergence rate, and juvenile mortality rate. The models developed will be useful for modeling and estimating the number of *Aedes aegypti* in the environment under different temperature, diurnal temperature variations, and larval densities.

**Author Summary:** Using experimental data from experiments conducted on *Aedes aegypti*, we formulated regression models to predict pupation, adult emergence, and juvenile mortality rates based on average temperature, temperature fluctuation range, and larval density. The prediction models produced were shown to account for high levels of variation within the data. Validation was performed by comparing omitted data sets to the predictions generated by our models. Our results show that the models produce results that are not significantly different from the experimental results and are capable of predicting aquatic development rates of *Ae. aegypti*.

## Introduction

The global burden of mosquito-borne pathogens such as dengue, chikungunya, and Zika virus, has been increasing with changing climate and the expansion of mosquito populations into new areas [1–3]. *Aedes aegypti* are well known to be competent vectors of these pathogens and flourish under warm and temperature climates [4–8]. Feeding primarily on humans and breeding in man-made containers in close proximity, *Aedes aegypti* are among the species of mosquitoes that are contributing to the dispersion of mosquito-borne pathogens in human populations [9,10].

Rainfall and nutrients are necessary factors for the viability of juvenile mosquitoes to hatch and develop, however, several other environmental factors such as exposure to light, average temperature, and diurnal temperature fluctuation ranges have been shown to impact the rates at which juvenile mosquitoes develop [11–24]. Prior studies have shown that larval density within aquatic breeding sites also play a role in the rate at which larvae pupate and pupae emerge as adults [25–30].

In this study, we utilize results from experiments detailed by Zapletal et al. [30] and fit a regression model to predict the rates of *Ae. aegypti* pupation, emergence and mortality under varying conditions of average temperature, diurnal temperature range, and larval density. Validation using additional experimental data was conducted to assess the validity of the model. Estimating the development rates of *Ae. aegypti* will provide greater insight into the population dynamics of this mosquito species across are variety of environmental conditions.

## Results

The average rate of pupation was positively correlated with average temperature (X_1_), temperature range (X_2_), and larval density (X_3_). Insignificant variables were eliminated from the model (α < 0.01) and only significant variables were retained in the model. The predictive model structure was identical for average pupation, average emergence, and average mortality rates. The model structure is provided in Equation 1, with the coefficient for each predictor provided for each model in Table 1 and full analysis of each predictor provided in Tables S1-S3.

*Equation 1. Predictive model structure*

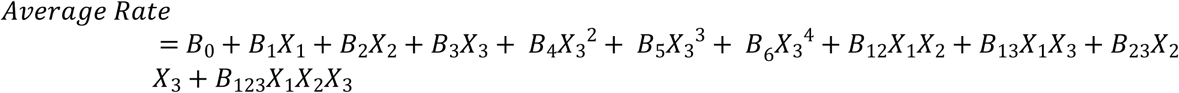

**Table 1.**
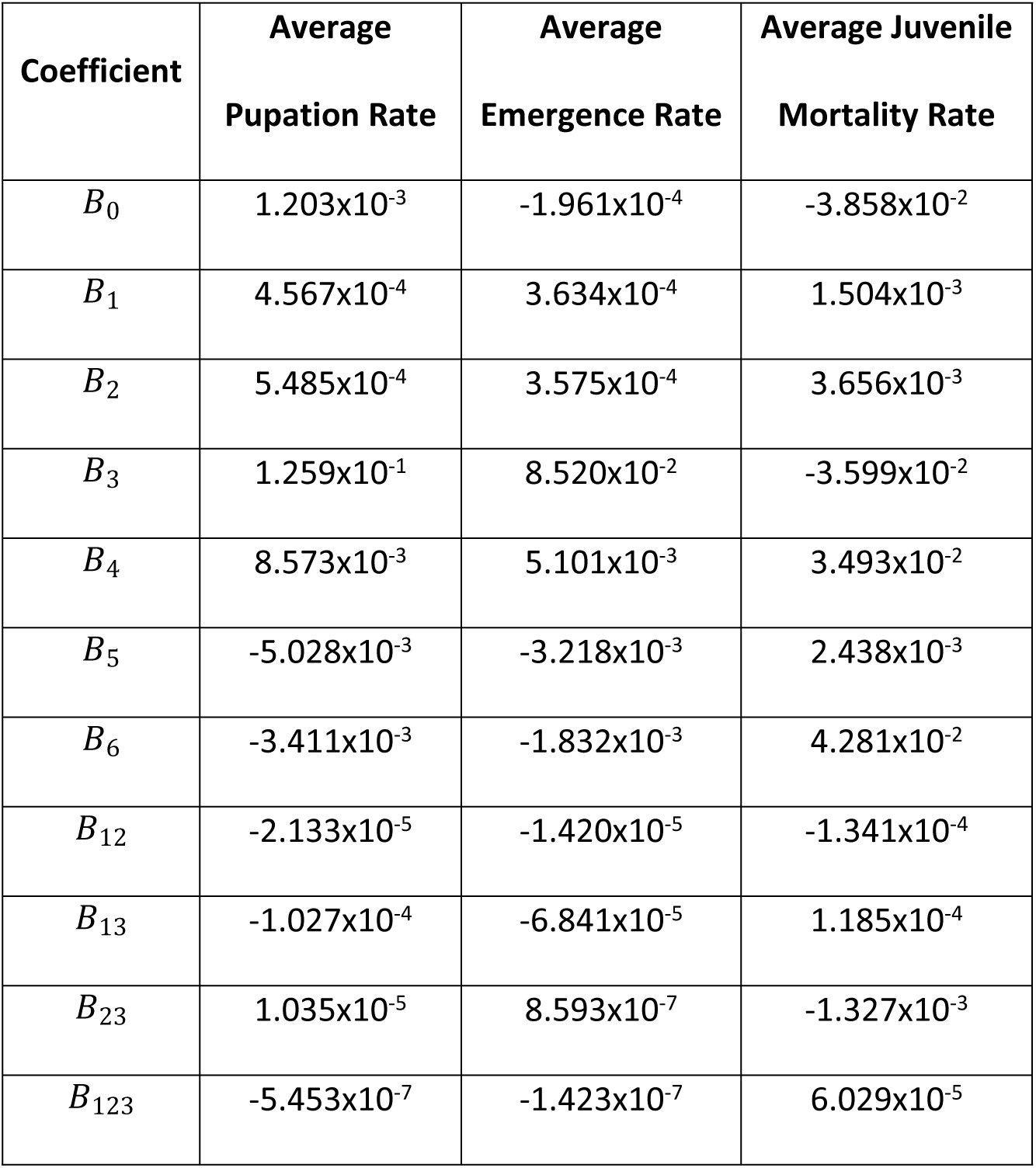
Predictor estimates for average pupation, adult emergence, and juvenile mortality rates

The prediction model for average pupation rate was consistent with experimental observations. The model showed a positive correlation between average temperature and average pupation rate, demonstrating that increases in average temperature increased the rate of pupation. Likewise, a positive correlation of temperature range showed that as temperature fluctuation increased an increased average pupation rate was observed. Larval density showed a positive correlation for degree one and two terms, but a negative correlation for degree three and four terms. As observed within the experiments, slower rates of pupation were associated with high larval densities. Two-factor interactions between average temperature and temperature fluctuation range, average temperature and larval density, and the three-factor interaction of all predictors were negatively correlated with average pupation rate. A positive correlation with average pupation rate was found for the interaction of temperature fluctuation range and larval density. Using all predictors and their interactions, the prediction model for average pupation rate produced an adjusted R^2^ of 0.8236. Paired t-test results (provided in Table S4) comparing the average predicted and observed pupation rates from excluded combinations provided a p-value of 0.731, showing no significant difference between the average predicted and observed pupation rates.

Similar to the model for average pupation rate, the prediction model for average adult emergence rate showed comparable results. With positive correlations of average temperature and temperature fluctuation range with average emergence rate, increases average temperature and temperature fluctuation range resulted in increases in average adult emergence rate. Furthermore, the larval density predictor showed positive correlation for first and second degree terms, but showed a negative correlation for third and fourth degree terms. Hence, as larval density increased, average adult emergence rate decreased. Two-factor interactions between average temperature and temperature fluctuation range, average temperature and larval density, and the three-factor interaction of all predictors were negatively correlated with average adult emergence rate. Only the interaction between temperature fluctuation range and larval density was positively correlated with average adult emergence rate. The prediction model for average adult emergence rate produced an adjusted R^2^ of 0.8577 and paired t-test results (provided in Table S5) show no significant difference between predicted and observe adult emergence rates with a p-value of 0.995.

Average juvenile mosquito mortality rates were positively correlated to average temperature and temperature fluctuation range, resulting in increased mortality rates as average temperature and temperature fluctuation rate increased. First degree terms for larval density were negatively correlated with average juvenile mortality rates, however, second, third, and fourth degree terms were positively correlated. This result agrees with what was observed during the experiments, as high levels of mortality were observed as larval density was increased. Two-factor interactions between average temperature and temperature fluctuation range, and temperature fluctuation range and larval density were shown to be negatively correlated with the average juvenile mortality rate. The two-factor interaction between average temperature and larval density, along with the three-factor interaction of all predictors were found to be positively correlated. The prediction model for average juvenile mortality rate produced an adjusted R^2^ of 0.7412. Results from the paired t-test (provided in Table S6) show no significant difference between predicted and observed mortality rates with a p-value of 0.535.

## Discussion

The results show that the models developed are adequate to predict the aquatic development rates of *Ae. aegypti* based on average temperature, temperature fluctuation, and larval density. The models of pupation, adult emergence, and juvenile mortality rates were validated by testing the models with predictor combination data excluded from the initial model training. Future laboratory experiments will focus on various average temperatures, larval densities, and smaller daily fluctuations of temperature to extend model coverage to micro fluctuations in diurnal temperature ranges. While the model predictions were consistent with the experimental observations, future experiments will increase model accuracy when applied to predictor combinations outside of those tested experimentally.

Although the prediction models showed no significant difference when compared to the experimental data, there were several limitations to this study. Only two average temperatures and two temperature fluctuation ranges were tested experimentally. While a linear relationship between these predictors is expected, future experiments will be necessary to confirm this pattern. Extension of the predictor models outside of the predictor ranges tested also poses a challenge. It has been demonstrated that mosquito aquatic development slows at both lower thresholds of 16°C and upper thresholds of 37°C, with fluctuations in temperature further impacting the development rates [31]. Additionally, a non-linear relationship between larval density and development rate may be exist beyond the larval densities examined. This may be, in part, due to the fact that during the larval stages of development, larvae excrete juvenile hormone that slows the development of the other larvae in breeding container. Hence, as the density of larvae increases, the concentration of juvenile hormone also increases. This concentration can reach a point where there will be no pupation from any of the larvae in a container. As a result, higher larval densities are potentially approaching a pupation rate asymptote of 0 pupae per hour. Similarly, since the emergence rate depends on the number of larvae that have pupated, the average rate of emergence will follow a similar trend. Average mortality rate will follow an inverse relationship, increasing with larval density and reaching a maximum rate at a threshold density. Further experiments over wider density ranges are required to identify the threshold value and maximum mortality rate.

The experiments presented in Zapletal et al. were conducted under optimal conditions of nutrition for mosquito development with abundant food supply and competition-reducing conditions. Thus the developments rates derived in this work provide an upper bound to the development rates in the field under sub-optimal nutrition availability conditions. Collection of data on micro-climatic conditions showed that the conditions under which mosquitoes develop can vary significantly from the ambient data collected in the environment [32,33]. The preliminary results have shown that temperatures within these micro-climates can be significantly warmer during winter months, allowing mosquitoes to survive even with ambient temperatures falling below 0°C.

Estimating the impact of different environmental conditions on *Ae. aegypti* will allow for a better understanding of mosquito population dynamics and provide a more accurate prediction of mosquito population. Furthermore, models that can capture these dynamics at a container level, relying on the individual characteristics and micro-climatic conditions at individual breeding sites, will allow for more accurate models of mosquito populations. The models presented in this paper will provide mosquito development rates at different environmental conditions for use in such detailed mosquito population models. Our future effort is focused on developing such a granular mosquito population model that will leverage the development rates derived from the models presented in this paper.

## Conclusion

With increasing global climate, the number of people impacted by *Ae. aegypti* and the diseases spread by this vector is increasing [34–37]. Prior work has been conducted to understand the impacts of environmental conditions that play a role in the development of these mosquitoes. This study explored the development of regression models to the aquatic development of *Ae. aegypti* by using the results from the experiment conducted by Zapletal et al. [30]. Regression models for rates of pupation, emergence, and mortality were developed using average temperature, temperature fluctuation range, and larval density as predictor variables.

Additional experiments following the same protocol as the prior experiments were conducted to obtain data points for validation. The validation experiments were designed to test an average temperature and larval densities that were not used to build the regression model. The comparison between the predicted and observed values showed that the model was sufficient to predict the average rates of pupation, emergence, and mortality. While the results showed reasonable errors, further experiments are necessary for the validation of temperature fluctuation range, as well as the upper bounds for larval density.

These regression models will prove to be useful in future modeling efforts to more accurately estimate mosquito populations. While previous mosquito population models have been developed, many of these models have overlooked the importance of larval density in the development rates of juvenile mosquitoes [38–43]. Also, previous work focused on developing population models at specific average temperature and specific diurnal temperature range. The regression models presented in this paper provide the flexibility to perform population analysis at any setting of average temperature, diurnal temperature range and density (within the bounds of ranges covered in the experiments). Our future research will utilize the prediction models of juvenile *Ae. aegypti* development and environmental data to examine impacts of micro-climatic conditions on mosquito population dynamics and the diseases that they spread throughout human population.

## Methods

Experimental data was obtained from previously published results in Zapletal et al. [30]. The experiments performed examined various aspects of aquatic *Ae. aegypti* development including time to first pupation, time of first emergence, maximum rate of pupation, time of maximum rate of pupation, maximum rate of emergence, time of maximum rate of emergence, final average proportion of adult emergence, and average proportion of larval mortality. The experiments conducted observed aquatic development rates under five larval densities (0.2, 1, 2, 4, and 5 larvae/mL of water), two constant temperatures (26.5°C and 32°C), and two diurnal temperature ranges (21-32°C and 26.5-37.5°C). Diurnal temperatures followed the daily temperature patterns, increasing linearly for nine hours from the low temperature in the morning (07:00) to the high temperature in the late afternoon (16:00) and then decreasing to the low temperature for 15 hours.

Using the data provided, a weighted average of the time to pupation and time to emergence were calculated according to the number of pupae or newly emerged adults recorded at each observation time. The average hourly rate of pupation and emergence was computed by taking the reciprocal of the average time of pupation and emergence, respectively. The hourly juvenile mosquito mortality rate was computed by summing the numbers of dead larvae and pupae and dividing by the total duration of the experiment (in hours). Three predictors were used to fit each response variable: (1) average temperature (26.5 or 32°C), (2) temperature fluctuation (0°C for constant or 11°C for diurnal temperature patterns), and (3) larval density (0.2, 1, 2, 4, or 5 larvae/mL of water). Linear regression models for average pupation, emergence, and mortality rates were created using Python 3 in Jupyter. To capture the non-linear effects, polynomial transformation of larval density (up to degree 4), interactions between average temperature, temperature range, and larval density were included in the model.

Five of the 20 predictor combinations of average temperature, temperature fluctuation, and larval density were excluded from the model training data and used for validation of the model predictions. These five combinations were excluded randomly, to ensure that each predictor value was omitted at least once. The excluded combinations are provided in Table 2. Predictions for each excluded combination were calculated and tested against the experimental values obtained by using a paired t-test. Statistical analysis was conducted using Minitab 17.

**Table 2.**
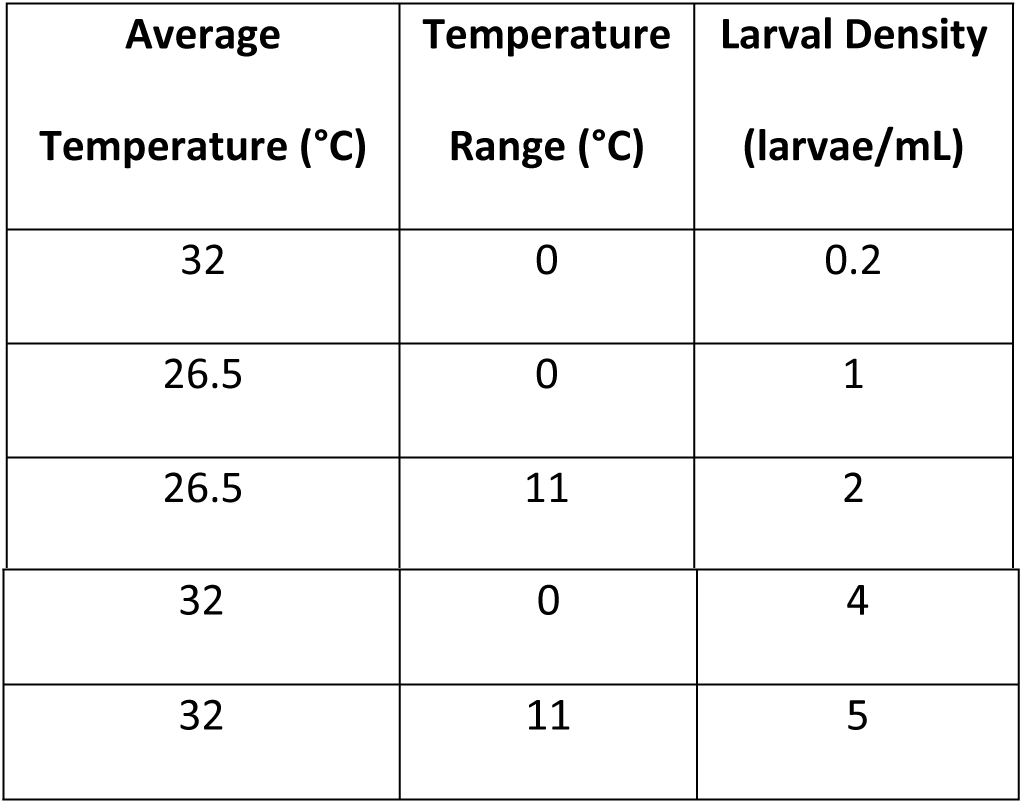
Excluded predictor combinations used for model validation

## Supporting Information

S1. Table. Analysis of predictor estimates and paired t-test results

